# A computational grid-to-place-cell transformation model indicates a synaptic driver of place cell impairment in early-stage Alzheimer’s Disease

**DOI:** 10.1101/2020.10.08.330928

**Authors:** Natalie Ness, Simon R. Schultz

## Abstract

Alzheimer’s Disease (AD) is characterized by progressive neurodegeneration and cognitive impairment. Synaptic dysfunction is an established early symptom, which correlates strongly with cognitive decline, and is hypothesised to mediate the diverse neuronal network abnormalities observed in AD. However, how synaptic dysfunction contributes to network pathology and cognitive impairment in AD remains elusive. Here, we present a grid-cell-to-place-cell transformation model of long-term CA1 place cell dynamics to interrogate the effect of synaptic loss on network function and environmental representation. Synapse loss modelled after experimental observations in the APP/PS1 mouse model was found to induce firing rate alterations and place cell abnormalities that have previously been observed in AD mouse models, including enlarged place fields and lower across-session stability of place fields. Our results support the hypothesis that synaptic dysfunction underlies cognitive deficits, and demonstrate how impaired environmental representation may arise in the early stages of AD. We further propose that dysfunction of excitatory and inhibitory inputs to CA1 pyramidal cells may cause distinct impairments in place cell function, namely reduced stability and place map resolution.

**Author Summary:** Cognitive decline in Alzheimer’s Disease (AD) correlates most strongly with dysfunction and loss of synapses in affected brain regions. While synaptic dysfunction is a well-established early symptom of AD, how impaired synaptic transmission may lead to progressive cognitive decline, remains subject to active research. In this study, we examine the effect of synapse loss on neuronal network function using a computational model of place cells in the hippocampal network. Place cells encode a cognitive map of an animal’s environment, enabling navigation and spatial memory. This provides a useful indicator of cognitive function, as place cell function is well characterized and abnormalities in place cell firing have been shown to underlie navigational deficits in rodents. We find that synapse loss in our network is sufficient to produce progressive impairments in place cell function, which resemble those observed in mouse models of the disease, supporting the hypothesis that synaptic dysfunction may underlie the cognitive impairment in AD. Furthermore, we observe that loss of excitatory and inhibitory synapses produce distinct spatial impairments. Future experiments investigating the relative contribution of different synaptic inputs may thus allow new insights into the neuronal network alterations in AD and potentially enable the identification of new therapeutic targets.

## Introduction

AD is the most common cause of dementia, marked by progressive memory impairment and other cognitive deficits that ultimately lead to a loss of functional independence [1]. The neurodegeneration in AD is associated with a reduction in neuronal activity and neuronal atrophy [1–3]. However, in patients and mouse models of AD, cognitive impairment correlates most strongly with synaptic dysfunction, which precedes cellular loss, suggesting AD is primarily a synaptopathology [4–8]. Furthermore, pathological forms of amyloid *β* and tau protein, key hallmarks of AD, have been found to trigger numerous molecular cascades that induce synaptic dysfunction and loss [4, 9, 10].

While it is well established that AD reduces neuronal activity, clinical observations, including the occurrence of epileptic seizures, suggest a more complex derangement of activity [11]. Recent neuroimaging studies have revealed that hyperactivity is the primary neuronal dysfunction in early AD, and may further exacerbate disease progression by increasing the release of amyloid *β* and tau protein [12–14]. There is substantial evidence suggesting an underlying disturbance of the excitation-inhibition balance as a result of impaired synaptic transmission [15]. For instance, impairments in slow-wave propagation in an APP/PS1 mouse model could be rescued by enhancing GABAergic inhibition [16]. An enhanced understanding of neuronal network alterations and the underlying actors may enable the identification of new therapeutic targets, and subtle changes in hippocampal activity provide promising early and easily-accessible biomarkers for diagnosis [17–19].

Pathological lesions of AD are primarily found in the hippocampus and associated structures [1]. Correspondingly, declarative episodic memory and navigation, which involve processing in the hippocampus, are typically impaired early [20]. Navigation and spatial memory require the activity of place cells, which encode a ‘cognitive map’ of an animal’s environment [21], and spatial impairments correlate with abnormalities in place cell activity in mouse models of AD [22–25]. Further experiments examining place cell dysfunction in early AD may shed light on how hippocampal network dysregulation is related to cognitive deficits. However, this is a challenging question to address experimentally, as it requires longitudinal tracking of individual cells and synaptic transmission. Using optical imaging, registration of the same neurons becomes increasingly difficult as the number of and interval between sessions increase [26]. Computational modelling may enable identification of critical timepoints and functional links to refine such *in vivo* experiments. The aim of this study therefore was to create a functional model of long-term place cell dynamics in hippocampal region CA1, in order to investigate how AD-related changes in connectivity impair network function throughout disease progression.

The computational model we have used here is based on a feedforward grid-cell-to-place-cell transformation, previously used with Hebbian learning and competitive network interactions to produce place field formation [27–30]. There is substantial evidence supporting the biological plausibility of grid cells as the major determinant of place cell firing. Grid cells are not only the most abundant type of spatially-modulated cell in the superficial entorhinal cortex (EC) [31], but also comprise a major proportion of the excitatory input to place cells [32]. In addition, place field size decreases in response to EC lesions [33], grid cell realignment coincides with place cell remapping [34] and grid cells have been implicated in the temporal organisation of place cell firing [35, 36]. In the absence of other inputs, direct EC input is sufficient to establish and maintain place cell firing [37], although spatial information content is reduced [38].

The model we present here is to our knowledge the first grid-cell-to-place-cell transformation model able to recapitulate the dynamics of CA1 place cell behaviour over long timescales, including long-term persistent place fields and stable place cell density [39–43]. Our modeling results provide strong support for the hypothesis that synaptic dysfunction drives cognitive impairment in early AD by disturbing the firing homeostasis of cortico-hippocampal circuits [15, 44, 45]. We further predict that excitatory and inhibitory synaptic dysfunction have distinct effects on place cell function in AD, suggesting a direction for future experimental work.

## Results

To study the effect of AD-related synapse loss on hippocampal place coding, we simulated a population of CA1 pyramidal cells, which received excitatory input from grid cells and inhibitory feedback from interneurons, beginning from the model of Acker et al. [28]. Place cell function was analysed in a 1 m linear environment, with cells characterised by their firing rates within each 1 cm bin along the track. The model was constrained to parameters that reproduced key characteristics of long-term CA1 place coding (see Methods). We then analysed and contrasted the effect of progressive excitatory and inhibitory synapse loss on network function to examine whether AD-related synapse loss may induce impairments in place cell function.

### Key requirements for long-term stability of CA1 place cell dynamics

In order to simulate place cell function over time, we implemented a grid-cell-to-place-cell transformation model previously described by Acker et al. [28]. In line with the recently established mean synapse lifetime of only 10 days in CA1 [46, 47], the model employed daily synapse turnover and demonstrated that Hebbian plasticity is sufficient to stabilize place fields, suggesting that memory can persist in the correlation structure of a dynamic network [48].

Our implementation of this model involved 10,000 grid cells and 2,000 pyramidal cells and produced dense place fields, with cells losing their place fields or remapping over time, while a subset of cells retained stable place fields (Fig 1A), as observed experimentally [42, 43]. However, the number of cells with significant place fields declined rapidly due to a lack of novel place field formation (Fig 1B), leaving only the subset of ‘long-term stable’ place cells. We therefore further constrained the model to those parameters that reproduced the stable place cell density observed experimentally [43, 49], which involved shorter feedback inhibition cycles, additional homeostatic constraints and a more realistic network architecture.

**Fig 1.**
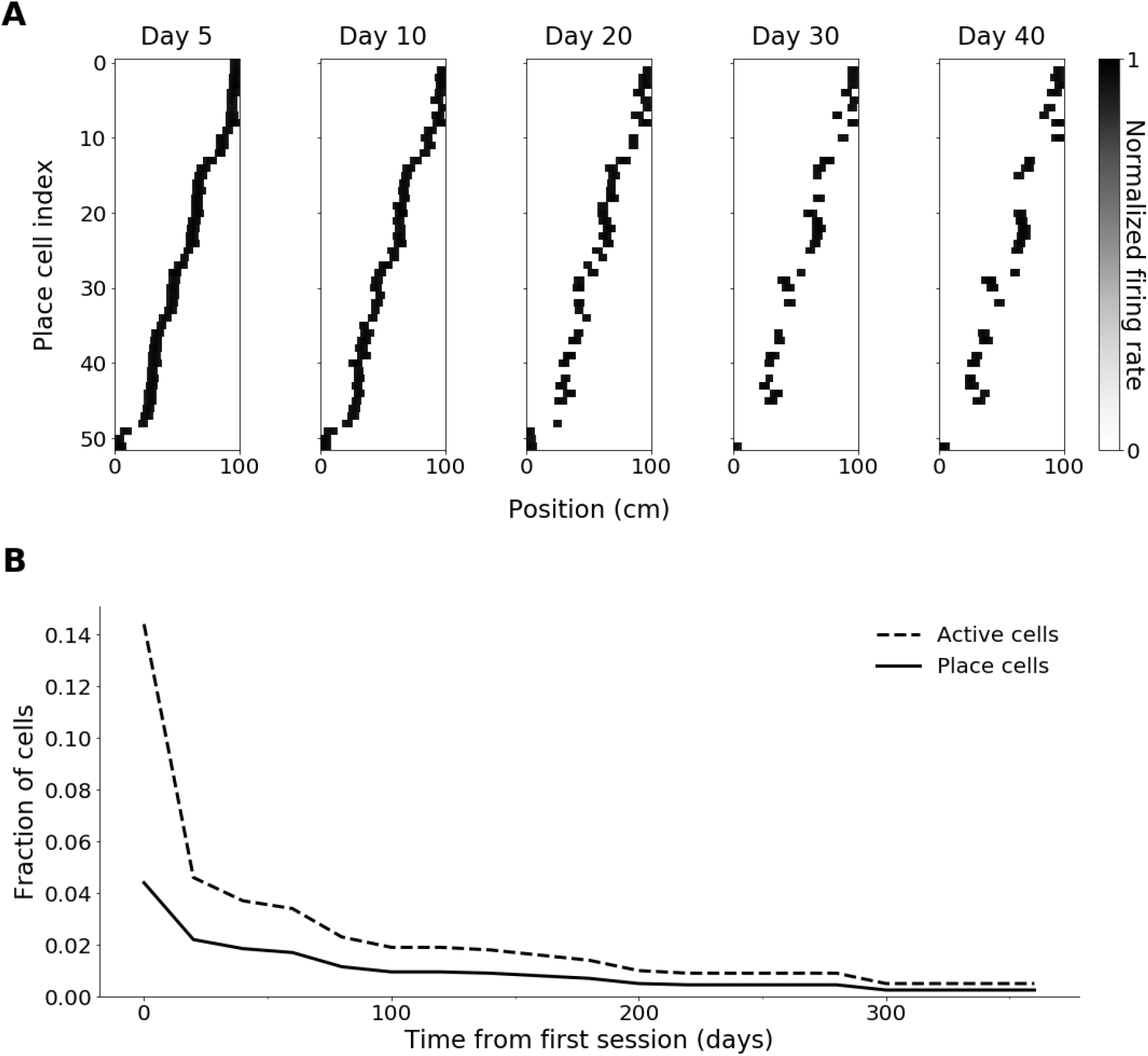
Simulation with long-term stability of place fields without novel place field formation. (A) Place field maps of cells with significant place fields on day 5, for days 5 to 40, indexed according to centroid position along the track on day 5. (B) Proportion of active cells and place cells over 365 days. Simulations were carried out with Hebbian learning.

### Improved feedback inhibition and synaptic plasticity

Interneuron-mediated feedback involved inhibition of pyramidal cells with firing rates below a given fraction of the highest firing rate. However, cell activities were compared across the entire track, whereas interneuron-mediated feedback is hypothesised to be precisely temporally coordinated, generating gamma-frequency rhythms [50, 51]. To better approximate this, feedback inhibition was restricted to individual bins along the track, which promoted firing in positions of low general activity with otherwise low place field coverage.

While synaptic scaling kept the average synaptic weight constant, the proportion of active cells decreased over time (Fig 1B). Strategies that aimed to counteract this instability by reducing competition included lower levels of Hebbian plasticity and constraints on synaptic weights and weight change, however, these approaches diminished stability, causing strongly fluctuating place cell numbers with no long-term stable cells.

To introduce additional homeostatic constraints, Hebbian learning was replaced by the Bienenstock-Cooper-Munro (BCM) learning rule [52]. BCM learning relies on basic Hebbian principles but incorporates a dynamic threshold for weight change that depends on recent postsynaptic activity and enables bidirectional modification of synapse strength. Consequently, high firing rates increase the threshold for potentiation and may result in synaptic depression instead, thus increasing competition. Under BCM learning, the rate of novel place field formation increased slightly, but not sufficiently, to balance place field loss.

### Unsuccessful strategies to stabilize place field density

As most forms of synaptic homeostasis observed experimentally operate over hours or days [53, 54], the rate of homeostasis was reduced. Multiplicative synaptic scaling does not allow for the homeostasis rate to be adjusted, so subtractive normalization was implemented. Under subtractive normalization, all synapses converging onto a neuron are changed by the same magnitude to keep the total synaptic input fixed, rather than weight adjustments being proportional to individual strength [55], thereby generating more competition [56]. Here, the sum of weights converging onto each cell rarely stabilized at the homeostatic point, as this would have required negative weights, yet the subtractive decay term was not sufficient to effectively limit growth of relatively stronger weights. Thus, subtractive normalization did not elicit the desired weight stabilization.

Incorporation of multiple enclosures was motivated by theories that place fields from different enclosures may ‘mix’ in familiar environments [57]. Running sessions in novel environments, represented by distinct grid cell arrangements [34], induced new place fields that were retained in the familiar environment through their effects on synaptic weights. However, continuous introduction of new enclosures was required to prevent stabilization at a specific subset of place cells, but still resulted in a general decline with occasional spikes in place cell number. These spikes were likely mediated by close correspondence of the novel and familiar grid inputs, facilitating rapid place field formation.

It has also been proposed that the co-existence of more and less stable stimulus responses may result from a diversity in synaptic plasticity [58]. Indeed, applying diverse learning rates across cells resulted in more place field formation than the BCM rule alone, however, again, it was not sufficiently high for stabilization. Furthermore, there was a stronger afferent synaptic strength divide, such that cells with slow plasticity were generally outcompeted.

### Stable place cell density through improved connectivity

The CA1 network was scaled according to relative cell and synapse numbers in the rat [59](Fig 2A), as there was less extensive data available for the mouse. Layer III of the EC is estimated to consist of 250,000 principal cells, 80% of which are spatially tuned [60, 61]. Using estimated CA1 population sizes [59], this gives a pyramidal to grid cell ratio of 1.5575:1 and a pyramidal cell to interneuron ratio of 8:1. Due to the vast diversity and incomplete functional characterisation of interneurons, a single hypothetical type of ‘average’ interneuron was used. For simplicity, synaptic connections among pyramidal cells and interneurons were omitted.

**Fig 2.**
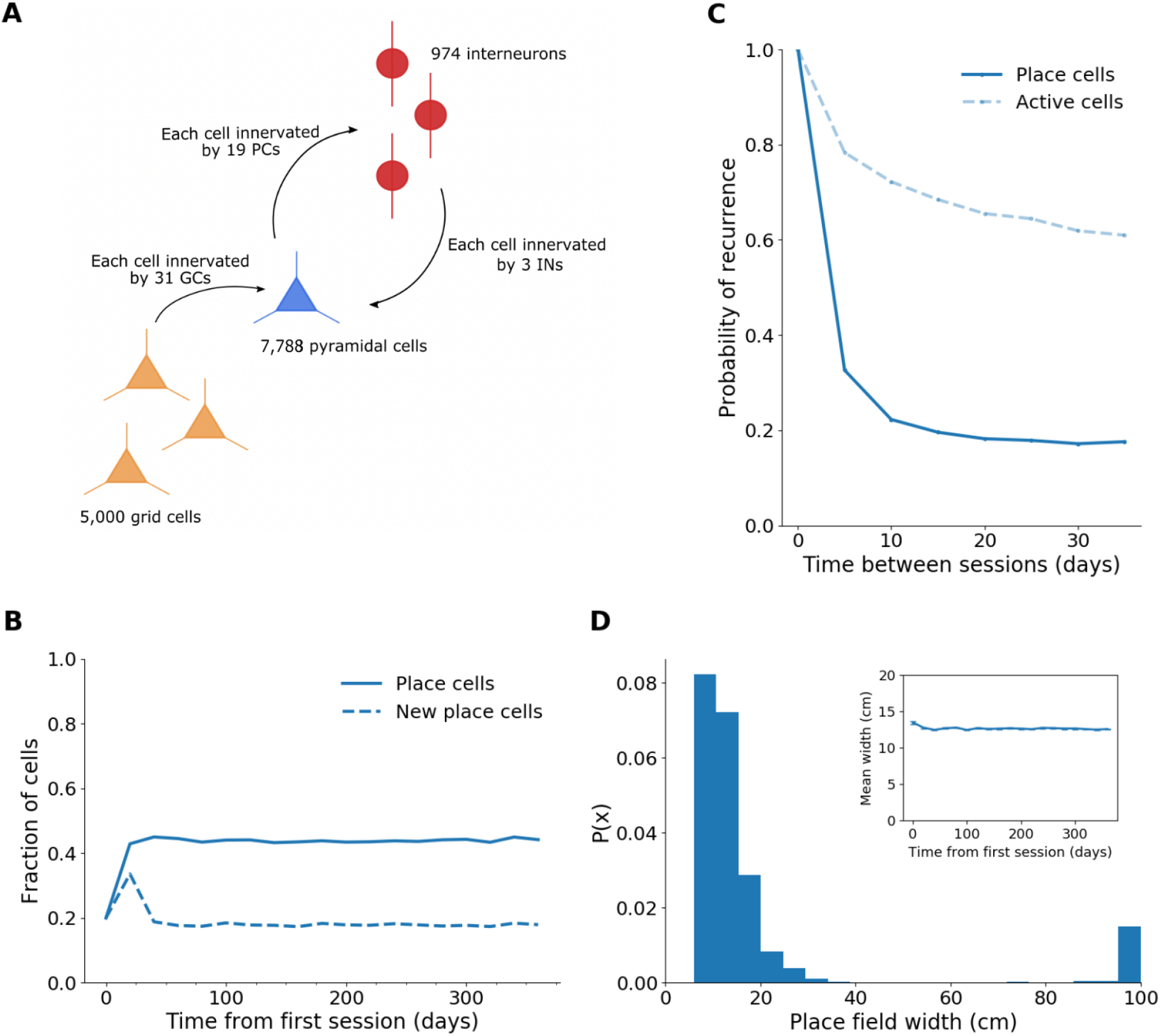
Stable place cell density and properties in the scaled model. (A) Schematic of new CA1 connectivity. (B) Proportion of total and new place cells over 365 days. (C) Probability of recurrence for active cells and place cells across sessions 5 to 35 days apart. (D) Representative probability density function of place field width. Inset: Mean place field width over 365 days, excluding widths > 50 cm. Error bars show standard error. Simulations were carried out with BCM learning.

As a result of lower excitatory synaptic density, pyramidal cell firing rates were lower but remained stable. The mean firing rate over 365 days was 0.52 Hz, with a range of 0-8.23 Hz, which corresponds closely to mean firing rates detected previously in CA1 place cells *in vivo* [62–65]. Interestingly, the network reproduced the stable place cell density observed experimentally, with cells dynamically losing and gaining place fields (Fig 2B-C). On average, 61% of cells were active each day, 72% of which formed significant place fields. Place field width ranged between 5 – 40 cm with a mean width of 12.6 cm (Fig 2D). As some cells fired throughout most of the track but there were no place fields of intermediate width (Fig 2D), cells with firing fields of > 50 cm were not classified as place cells.

Stable place field emergence in this model suggests the previous high synaptic density may have been a limiting factor by restricting variability in cell input even in the presence of high synaptic turnover.

The probability of recurrence of place cells with place fields that have drifted less than 5 cm between any two sessions, a measure of how likely it is for a cell to retain its place field, decreased from 98% and 51.9% for sessions 5 and 30 days apart, respectively, to 32% and 17.2% in the new model (Fig 2B). At stable place cell density, this indicates increased variability in the place cell ensemble.

At sufficient network size, determined to be approximately 5,000 grid cells and 7,788 pyramidal cells, the general features of the model are robust to cell number changes. However, as the model is scaled up, competition increases, leading to a lower proportion of place cells (10,000 grid cells: mean ± SEM = 19.9% ± 0.2%; 7,500 grid cells: 28.5% ± 0.1%; 5,000 grid cells: 43.8% ± 0.1%; *F* (2, 18) = 7164.82, *p* < 7.6^−27^).

The general features are also robust to the choice of inhibition model, although using an ‘E%-max’ inhibition model, in which cells whose firing rate is within 10% of the most excited cell escape inhibition, as opposed to a ‘winner-takes-all’ model, increases the proportion of place cells due to lower competition (winner-takes-all: mean ± SEM = 29.0% ± 0.2%; E%-max: 43.8% ± 0.1%). The ‘E%-max’ model was used here as it accounts for feedback delay [66].

### Excitatory synapse loss reduced firing and place map stability

In APP/PS1 mouse models, progressive reductions in spine density on the distal dendrites of CA1 pyramidal neurons, which are innervated by EC layer III, have been detected at 4-15 months of age [67], with a similar extent of loss occurring in 5xFAD models [68].

This extent of grid-to-pyramidal cell synapse loss in our model gradually reduced the median firing rate, while increasing the range of firing rates (day 20: median ± std = 0.34 ± 0.72 Hz, range = 3.71 Hz; day 360: 0.26 ± 0.73 Hz, range = 4.45 Hz). In addition, the firing rates became more uniform with less strongly pronounced ‘bumps’ (Fig 3A-B).

**Fig 3.**
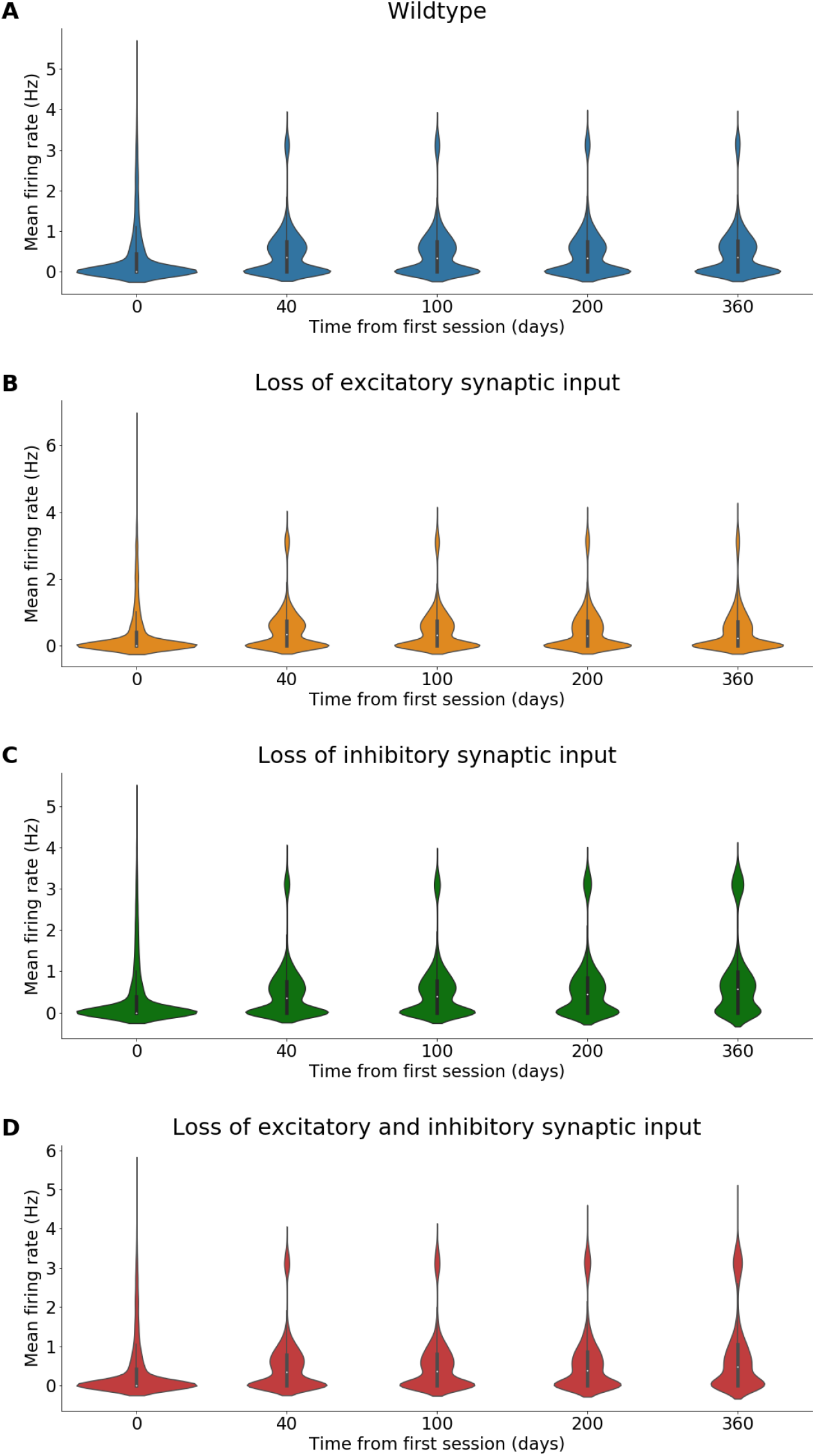
Each model exhibits characteristic changes in firing rate. Violin plots showing mean firing rates of CA1 pyramidal cells over 360 days in the wildtype (A), the grid-to-pyramidal-cell synapse loss model (B), the interneuron-to-pyramidal-cell synapse loss model (C) and the double synapse loss model (D). Median indicated by white dot within boxplot.

Synaptic scaling led to a compensatory increase in the mean and range of synaptic weights (Fig 4B), which likely mediated the relatively small reduction and increased variability of mean firing rates.

**Fig 4.**
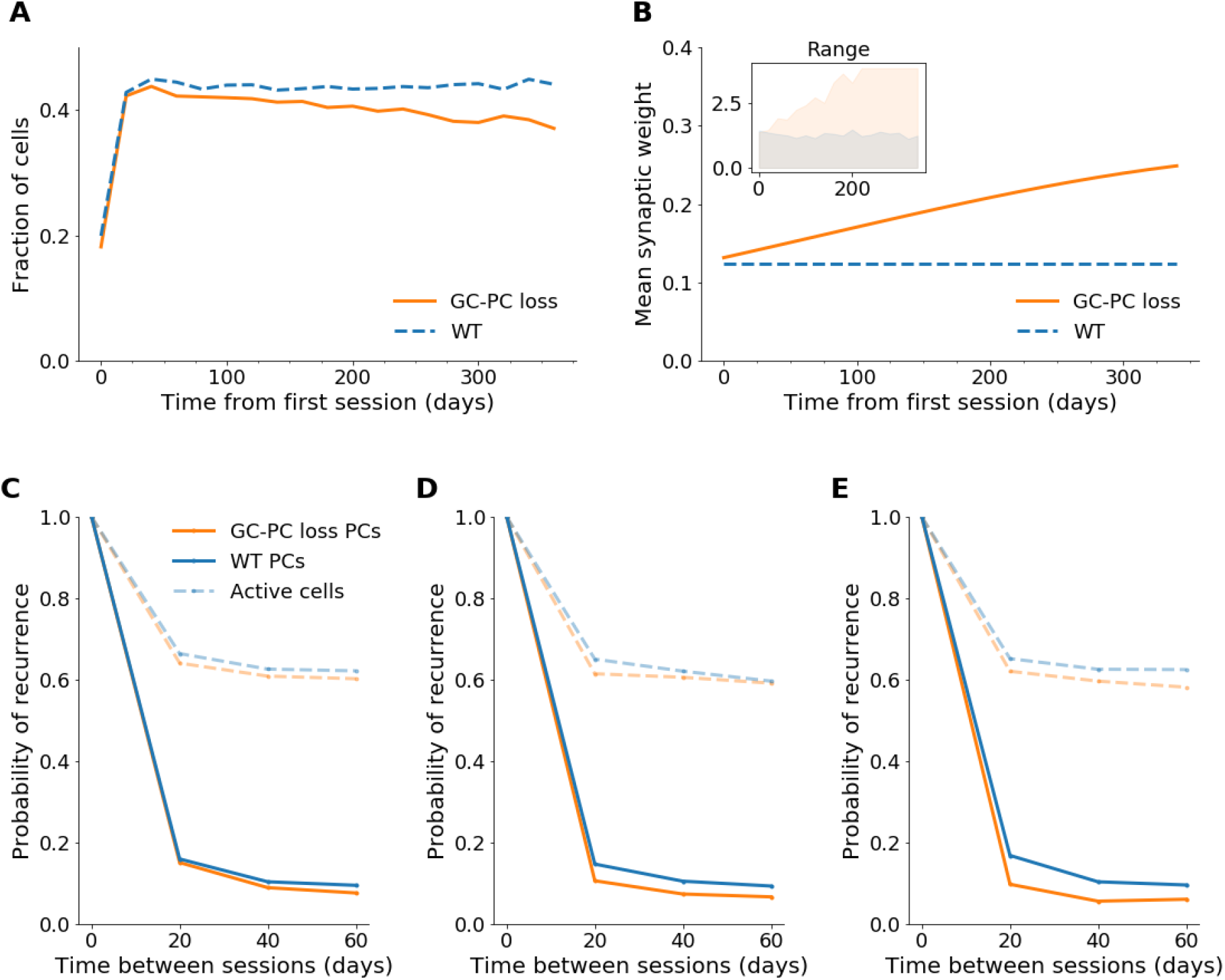
Place cell proportion and place map stability decreased as excitatory synapses were lost. (A) Proportion of place cells over 365 days in the grid-to-pyramidal-cell (GC-PC) synapse loss and wildtype (WT) model. (B) Mean synaptic weight over 365 days. Inset: Range of weights in the GC-PC loss model (orange) and the wildtype (blue). (C-E) Probability of recurrence for place cells and active cells across sessions 20 to 60 days apart from day 20 (C), day 200 (D) and day 300 (E).

The proportion of place cells decreased (Fig 4A), while the remaining cells retained a constant place field width (mean ± SEM = 12.6 ±0.02 cm). The reduced number of place cells may have slightly diminished coverage of the environment. In addition, the probability of recurrence of active cells and place cells between sessions 20 days apart, declined over time (Fig 4C-E), indicating lower stability.

### Inhibitory synapse loss increased neuronal activity and place field size

Progressive axonal loss, with normal bouton density on unaffected axons, has been detected on CA1 O-LM interneurons in 4-11 months old APP/PS1 mice in two separate studies [69, 70].

In our model, this extent of loss of inhibitory synapses caused a gradual increase in the median and standard deviation of mean firing rates (day 40: 0.35 ± 0.73 Hz; day 360: 0.58 ± 1.01 Hz), resulting in a significant difference to the wildtype (day 360: 0.35 ± 0.71 Hz; *p* < 4^−63^, Wilcoxon rank sum test) (Fig 3C). Furthermore, the fraction of place cells (Fig 5A) and active cells (day 20: 61.1%; day 360: 73.1%), as well as the probability that an active cell remained active between sessions (day 20: 65.4%; day 300: 72.5%) increased gradually. However, there was no difference in the recurrence probability of place cells between sessions 20 days apart (mean over days 20-300: 15.7%). Thus, impaired inhibition increased neuronal activity but did not affect across-session stability.

**Fig 5.**
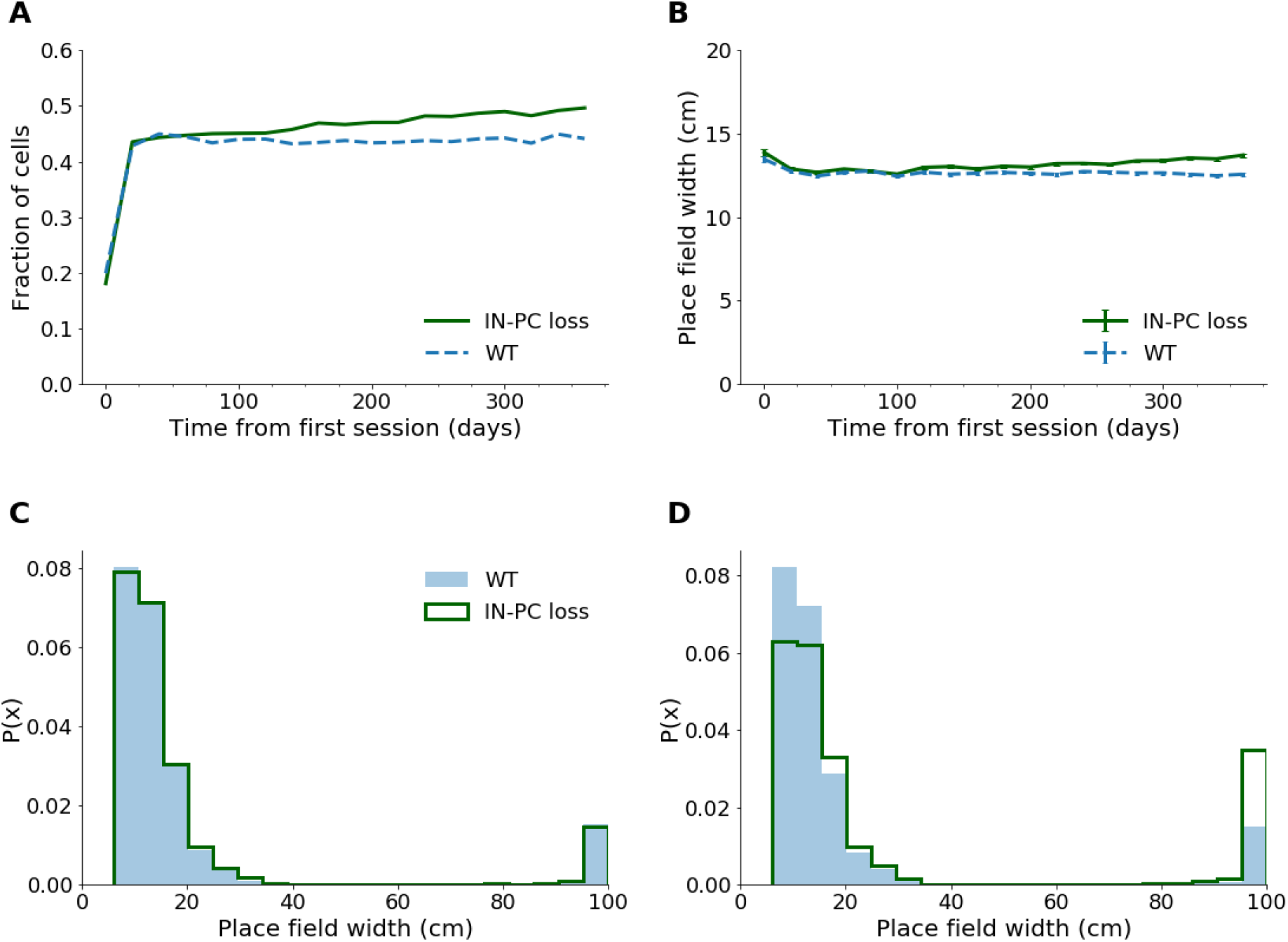
Place cell proportion and mean place field width increased over time when inhibitory synapses were lost. (A) Proportion of place cells over 365 days in the interneuron-to-pyramidal cell (IN-PC) synapse loss and wildtype (WT) model. (B) Mean place field width over 365 days, excluding widths > 50 cm. Error bars show standard error. Probability density function of place field widths on day 20 (C) and 360 (D).

Although the proportion of place cells increased, the fraction of active cells that form significant place fields decreased from 92.5% to 85.7% between days 20-360. This is likely linked to increased activity, as the proportion of cells with multiple place fields (mean ± SEM days 200-360: wildtype = 2.9% ± 0.09%; IN-PC loss: 3.6% ± 0.1%), which did not classify as place cells, and cells that were active throughout most of the track (Fig 5D) increased over time.

In addition to more place field formation, the mean place field width slightly increased (Fig 5B-D), indicating a reduced place map resolution.

### Excitatory and inhibitory synapse loss impaired place map resolution and stability

When both excitatory and inhibitory synapses were affected, the median neuronal activity also increased significantly (day 40: 0.35 ± 0.76 Hz, day 360: 0.49 ± 1.02 Hz; *p* < 1.1^−37^, Wilcoxon rank sum test) and the range of mean firing rates was higher than in the previous models (range of mean firing rates = 0 – 4.78 Hz; GC-PC loss: 0 – 4.04 Hz, IN-PC loss: 0 – 3.79 Hz)(Fig 3D).

In addition to a progressive increase in the proportion of highly active cells (Fig 3D), more frequent changes in the firing rate could be observed (Fig 6). For instance, in the wildtype just under half of all silent cells remained silent across sessions 20 days apart, while this was only true for a third of cells in the synapse loss model. Nevertheless, activity shifts were mostly gradual in both models, as exemplified by most new highly active cells coming from the pool of ‘intermediately active’ cells (Fig 6).

**Fig 6.**
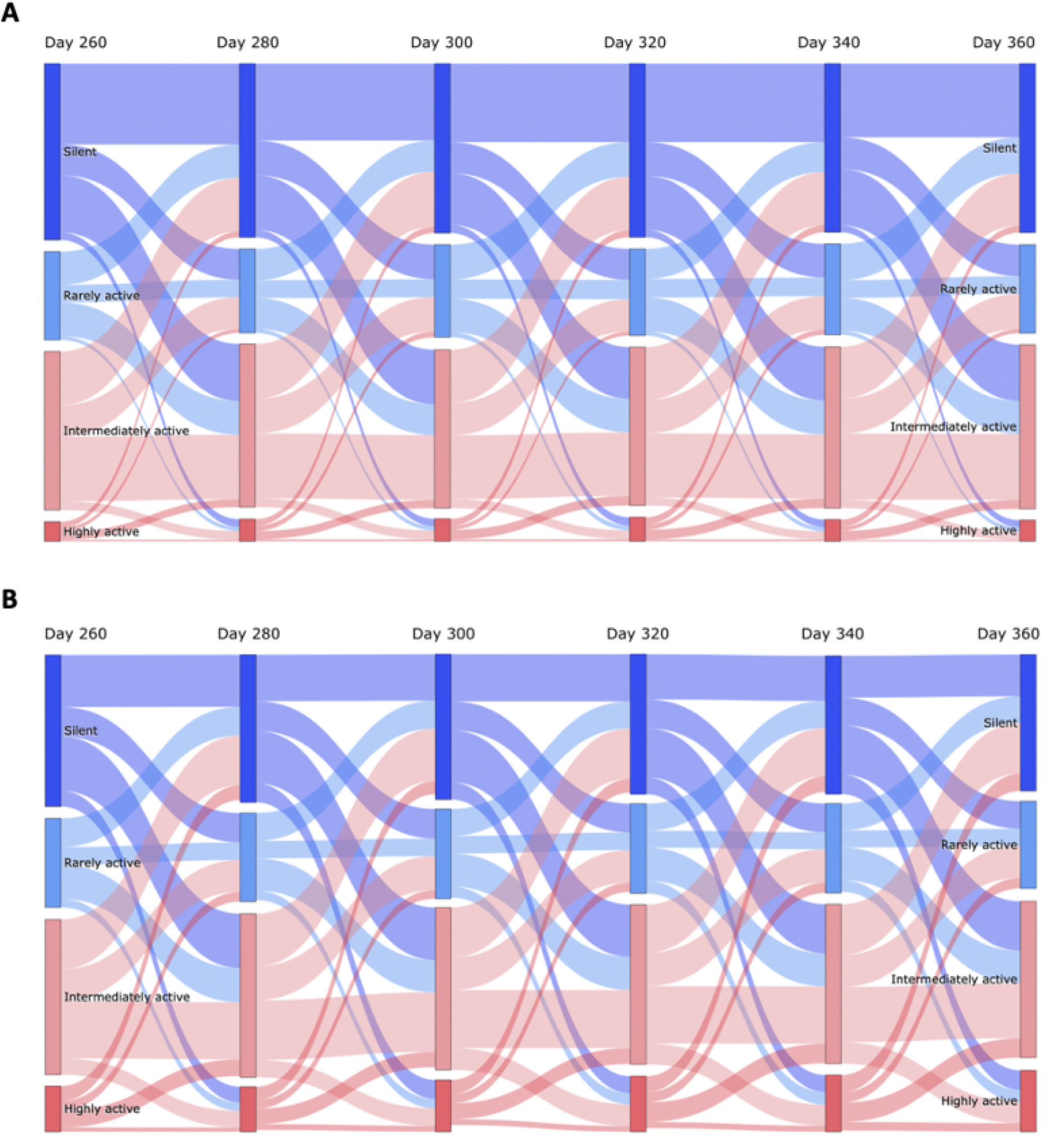
The proportion of highly active cells increases and firing rates are less stable in the synapse loss model. Sankey diagrams showing the proportion of cells classified by mean firing rate and flow between categories across sessions 20 days apart for the wildtype (A) and the synapse loss model (B) starting on day 260. Cells were classified as silent (mean firing rate = 0 Hz), rarely active (0 − 0.5 Hz), intermediately active (0.5 − 2 Hz) and highly active (> 2 Hz).

The proportion of place cells remained constant over time (mean ± SEM = 43.5% ± 0.01%)(Fig 7A), despite increased activity. There was a gradual increase in mean place field width (day 20: mean ± SEM = 12.7 ± 0.1 cm; day 360: 13.8 ± 0.1 cm)(Fig 7B) comparable to the inhibitory synapse loss model (day 360: 13.7 ± 0.1 cm). Furthermore, there was more variability and abnormal place field sizes (Fig 7B), likely mediated by a lack of inhibition in conjunction with lower excitatory input.

**Fig 7.**
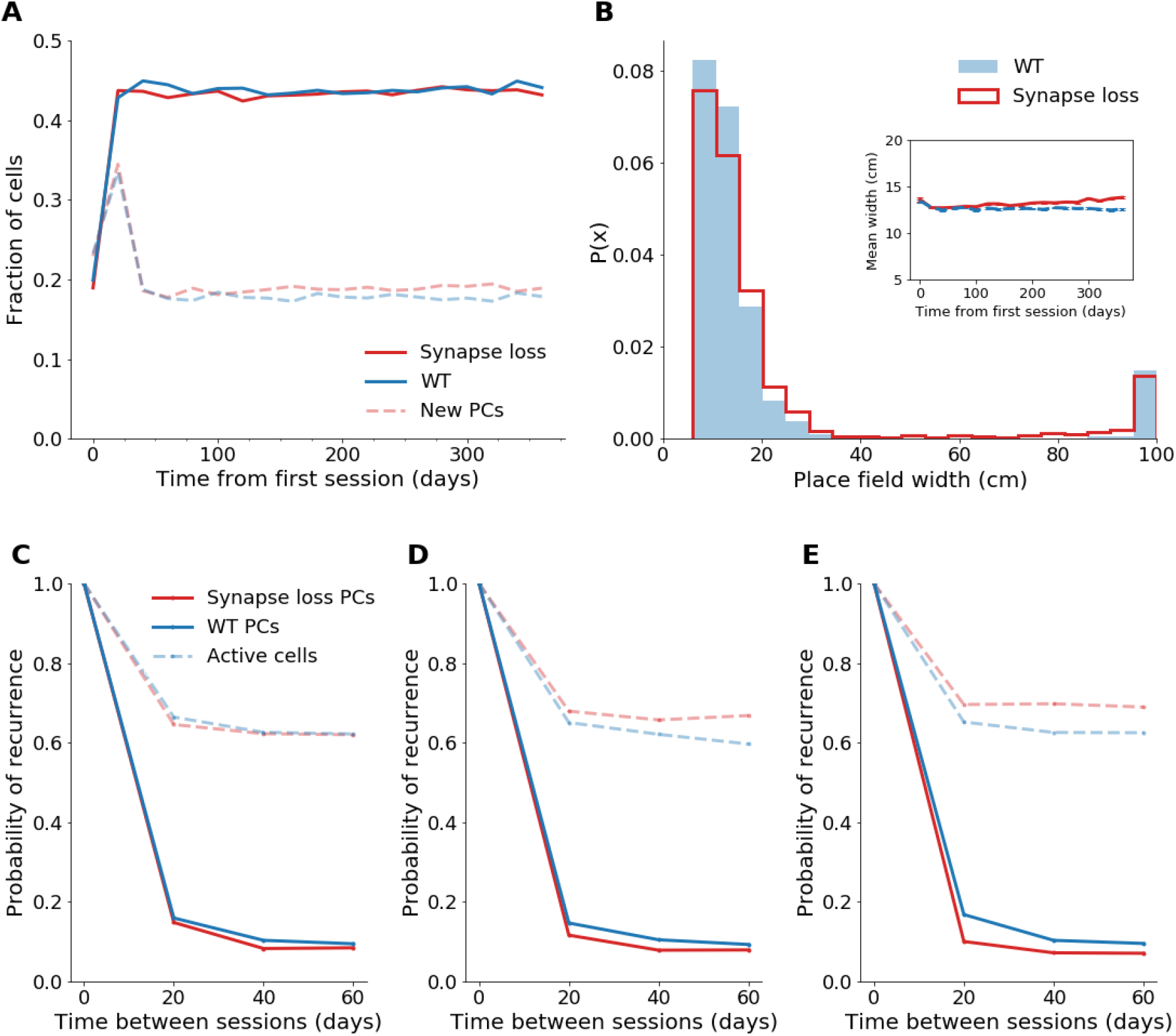
Place cell properties diverge from the wildtype in the double synapse loss model. (A) Proportion of total (dark lines) and new (pale lines) place cells over 365 days in the double synapse loss and wildtype (WT) model. (B) Probability density function of place field widths on day 360. Inset: Mean field width over 365 days, excluding widths 50 cm. Error bars show standard error. (C-E) Probability of recurrence for active cells and place cells across sessions 20 to 60 days apart from day 20 (C), day 200 (D) and day 300 (E).

As in the excitatory synapse loss model, the recurrence probability of place cells was gradually reduced (Fig 7C-E). Place cell density was maintained by a slight increase in new place cells (mean ± SEM for days 40-360: 18.8% ± 0.1%, wildtype: 17.8% ± 0.1%)(Fig. 7A).

## Discussion

In this study, we investigated the effect of AD-related synapse loss on hippocampal network function using a computational model, in which place fields were generated through the interplay between excitatory grid cell input and inhibitory feedback from interneurons. By incorporating homeostatic mechanisms and synaptic connectivity based on experimental findings, the model was able to replicate key aspects of CA1 place cell dynamics. To our knowledge, this is the first model that generates stable place cell dynamics from grid cell inputs long-term. By focusing on place cell function, the model can be used to predict how alterations at the synapse level may affect the emergent properties of the CA1 network.

### Development of the dynamic CA1 network model

To increase place field formation, interneuron-mediated feedback on more plausible timescales and BCM learning were implemented. While Hebbian plasticity is dominantly used in place cell models, the BCM rule has also been shown to support place coding [71, 72]. However, to enable temporal competition using the BCM rule, the rate of synaptic change must be substantially slower than the rate of input presentation, which requires extensive environmental sampling [52]. In contrast, computational modelling suggests fast synaptic changes are required to generate place fields at a realistic timescale [73]. This implies that place coding may involve a learning scheme other than the BCM rule, or multiple learning schemes depending on the input. Alternatively, de Almeida et al. [27] have proposed that place field formation may not rely on synaptic plasticity, except for refinements and long-term stability. They demonstrated place field formation in a grid-cell-to-place-cell model without learning, consistent with close to normal CA1 place coding during exploration in mice with non-functional N-methyl-D-aspartate-receptors (NMDARs), despite exhibiting an NMDAR-dependent form of long-term potentiation [74]. Furthermore, features of BCM learning have been observed in the hippocampus, most notably a dynamic postsynaptic-activity-dependent threshold for synaptic strengthening [75, 76]. As such, BCM-like learning remains a plausible alternative to Hebbian learning and has successfully produced the desired grid-cell-to-place-cell transformation at the timescale considered here.

Place cell density was stabilized using a more biologically plausible connectivity, thereby facilitating more realistic feedback inhibition and more dynamic grid cell input through lower synaptic density.

Our model also incorporated synaptic scaling, for which underlying biophysical mechanisms have been identified, including activity-dependent modulation of glutamate receptor expression [77]. While homeostatic mechanisms operate at a slower timescale *in vivo*, modelling studies commonly speed up homeostasis to stabilize Hebbian learning [78–80]. The mechanisms underlying this temporal contradiction between Hebbian plasticity and homeostasis remain unclear, however, proposed explanations include a yet unidentified rapid compensatory process [78].

In mice, it has been reported that 31% of CA1 pyramidal cells are active, while 20% form significant place fields in an environment [43]. While comparisons are difficult due to differences in experimental setup and the possibility of missing silent cells in *in vivo* recordings, our model, with 61.0% active cells and 43.8% place cells, likely involves a higher proportion of place cells than is observed *in vivo*, even compared to rat estimates [81, 82]. As the fraction of active cells and place cells decreased with increasing cell numbers, this may be due to the small network size.

Place map stability, with a recurrence probability of place cells of 32% and 17% for sessions 5 and 30 days apart, respectively, is comparable to experimental values of 25% and 15% [43]. As observed *in vivo*, the recurrence probability was found to only be moderately time dependent.

The mean place field size of 12.6 cm was relatively small, with reported lengths in 1m linear tracks ranging between 20 − 25 cm [43, 83]. Place fields are generally smaller in computational models [28, 29, 84] and it has been argued that grid cell inputs may not be sufficient to produce realistic field sizes without additional input from weakly spatially-modulated cells or recurrent connections among CA3 place cells [85]. In some studies, place field size has been increased by adjusting the grid scale [29, 84]. Thus, there are multiple strategies that may enable more realistic place field sizes, however, such discrepancies in spatial scale are unlikely to significantly affect the network properties [29].

Ziv et al. [43] have found that place cells generally maintain identical place fields *in vivo*, whereas our model exhibited a high level of remapping, prompting strict positional constraints on place fields considered ‘identical’ across sessions. The proportion of new cells within the place cell ensemble increased with cell number (7,788 pyramidal cells: 30.3%, 11,682: 37.9%, 15,575: 39.3%), indicating the involvement of a larger pool of cells. Thus, the frequent remapping may have been caused by the small network size.

### Excitatory synapse loss may reduce place map stability

Progressive loss of excitatory synapses caused synaptic weight rearrangements consistent with compensatory spine growth and glutamate receptor expression observed *in vivo* and in AD patients [67, 68]. Despite synaptic scaling, the median firing rate decreased and there was more firing variability. The decreased activity may have also impacted interneuron function by lowering the inhibition threshold, thereby providing an additional buffer for the firing rate. Lower mean firing rates in combination with some highly active neurons have also been observed in the CA1 region of 3xTg and APP/PS1 mice, in which this was accompanied by lower entropy, a measure of firing pattern diversity, indicating reduced coding capacity [86, 87].

There was a lower proportion of place cells compared to the wildtype, which has also been observed in 6-months-old APP/PS1 and APP-KI mice [23, 24]. Our model also suggests that excitatory synapse loss could mediate reduced stability of place maps. As reduced stability coincides with spatial memory impairments in APP-KI and 3xTg mice [23, 87], which also exhibit synaptic abnormalities [88, 89] but have normal place field sizes [23, 87], it may be a key mechanism contributing to memory deficits in AD.

### Inhibitory synapse loss may link hyperactivity to reduced map resolution

As interneurons play a key role in the spatiotemporal control of neuronal activity [15, 90], disturbances of which correlate strongly with cognitive deficits [24, 91, 92], the ‘GABAergic hypothesis’, which suggests impaired inhibition may be a critical link between the diverse dysfunctions occurring in AD, has gained popularity in recent years [15].

The increase in firing rate, proportion of highly active cells and cells that were active throughout the whole track is consistent with the hyperactivity observed in CA1 *in vivo* [93]. Place fields were larger and showed increased variability compared to the wildtype, as has been observed in Tg2576 and APP/TTA mice in conjunction with lower spatial information content [22, 25]. Increased place cell number and place field size indicate a lower resolution place map, as hypothesised to occur in experimental models [25].

### Excitatory and inhibitory synapse loss may distinctly contribute to network dysregulation

Korzhova et al. [94] recently identified a progressive increase in activity of intermediately active cells as the primary source of highly active neurons in the cortex of APP/PS1 mice, which was also observed in our model, supporting their hypothesis that network pathology may stem from stable aberrant activity of single cells.

When both excitatory and inhibitory synapses were affected, the resolution and stability of the place map were significantly diminished, accompanied by increased neuronal activity. This combination of dysfunctions has also been detected in an APP transgenic model [25], and in a mouse model overexpressing synaptojanin-1 (Synj1), a regulator of synaptic function, which has been implicated in AD [95, 96]. This provides further support for synaptic dysfunction underlying place cell abnormalities and suggests that both excitatory and inhibitory synaptic dysfunction contribute to place cell dysfunction in AD and produce distinct impairments in environmental representation.

### Implications for future work

The causative link established here between synaptic dysfunction and cognitive impairments is in line with current theories implicating impaired functional connectivity as the major driver of AD pathogenesis [15, 19, 45].

Based on our findings, we predict that hyperactivity in CA1 should coincide with place cell abnormalities and spatial impairments. Furthermore, we hypothesise that impaired spatiotemporal control of pyramidal cell firing by interneurons may be a major contributor to abnormal place field size. Similarly, decay of excitatory EC inputs may underlie reduced ensemble stability. An experimental study employing EC lesions supports our finding that place map stability is reduced in the absence of such inputs, however they also reported increased place field sizes [97]. Recent evidence suggests pyramidal cells in EC layer II also directly innervate CA1 interneurons and that synapses between these populations are selectively lost in Tg2576-APPswe mice [98]. The Tg2576-APPswe model had enlarged place fields, which could be rescued by optogenetic activation of the EC-interneuron synapses. As such, place field enlargement in response to EC lesions may have been mediated by impaired interneuron function.

The relative contribution of excitatory and inhibitory synapse dysfunction in AD is still unclear but may enable the identification of new therapeutic targets. Due to the high inter-connectivity within the hippocampal circuit, addressing this question experimentally requires precise silencing of synaptic transmission, however silencing with the required spatiotemporal control is challenging, potentially limiting the feasibility of such experiments [99].

### Potential extensions to the model

A biophysical model would enable examination of other interesting network properties observed in AD, such as hypersynchrony [100]. The model could also be further improved by including synaptic weights, synaptic homeostasis and an explicit firing rate for the interneurons, ideally also accounting for the diversity of interneuron types in CA1. Homeostatic inhibitory mechanisms, including altered receptor levels and GABAergic synaptic sprouting have been reported *in vivo* [101, 102]. Furthermore, it would be interesting to extend the model to incorporate place cell remapping, as this has been found to be impaired in AD [23].

A general limitation of grid-cell-to-place-cell feedforward networks is recent evidence for place-cell-to-grid-cell feedback, including the disappearance of grid cell firing in response to place cell inactivation, although the function of this feedback remains unclear [103]. Furthermore, recent experimental findings have challenged the view that grid cells are the major determinant of place field formation, including the formation of place fields after grid cell disruption [97, 104] and the earlier emergence of place cell firing during rat development [105, 106]. Input from subcortical structures, the pre- and parasubiculum and CA3 have all been found to contribute to the spatial tuning of CA1 place fields [30, 107–109]. Spatial input from CA3, for instance, is essential for rapid contextual learning in novel environments, temporal coding at the population level and memory retrieval [38, 110]. In light of this evidence, Bush et al. [30] have proposed that place cell firing is determined by a variety of sensory inputs, with grid cells providing complementary self-motion-derived input to support precise navigation. However, the relative contribution of different cell populations to CA1 place coding remains subject to active research and may even be dynamic, varying with condition or state [111].

While synaptic loss has been found to occur prior to the formation of amyloid plaques [112], plaque proximity-dependent loss has been reported in mouse models [113–115]. As such, it would also be interesting to explore the effect of concentrated areas of synaptic loss. In addition, shrinking cell population sizes, affecting interneurons as early as 6-to-12-months of age, have been detected *in vivo* [70, 116–118], and may contribute to network dysfunction.

Overall, our model shows that synapse loss in CA1 is sufficient to generate the network abnormalities observed in experimental models and AD patients, including hyperactivity [93, 100]. Furthermore, the resulting network abnormalities were shown to induce place cell dysfunction, which has been hypothesised to underlie spatial memory impairment in AD. Thus, our model shows synapse loss is sufficient to drive progressive network dysfunction and impaired spatial representation.

## Methods

### Grid cell simulation

We simulated a 1 metre linear track, with cells characterized by their firing rate in each 1 cm bin. Grid cells were simulated as described by Blair et al. [119] (Eq 1), such that their firing rate varied in a hexagonal grid with orientation, phase and offset randomly chosen from a uniform distribution,

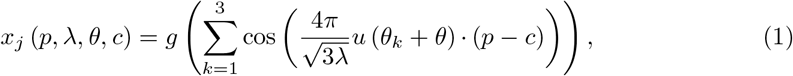

where *x_j_*(*p, λ, θ, c*) is the firing rate of grid cell *j* at position *p* with inter-vertex spacing *λ*, angular offset *θ* and spatial phase *c*. *λ* varies between 30 − 100 cm, *c* ranges between 0 − 100 cm and *θ* ranges between 0 − 60°. ‘·’ indicates the dot product operator. *θ_k_* is given by (cos (*θ_k_*), sin (*θ_k_*)), such that the cosine function gives a pattern of alternating maxima and minima in direction *θ_k_*. The hexagonal grid is generated by the sum of cosine patterns *θ*_1_ = −30°,*θ*_2_ = 30° and *θ*_3_ = 90°. *g* (*x*) = exp [*a* (*x − b*)] − 1 is a monotonically increasing gain function with *b* = −3/2 and *a* = 0.3, giving a firing rate of 0 − 3 Hz. Parameters were adapted from de Almeida et al. [27].

### Pyramidal cell simulation

Pyramidal cell activity was modelled as the sum of excitatory grid cell inputs [27],

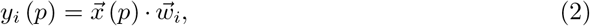

where *y_i_* (*p*) is the firing rate of pyramidal cell *i* at position *p* in Hz, 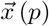 is the vector of firing rates of all grid cells that project to pyramidal cell *i*, at position *p*, and 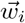 is the vector of synaptic weights converging onto pyramidal cell *i*.

### Synaptic weights

The initial weight *w_ij_*(*s*) of a synapse between grid cell *j* and pyramidal cell *i* was assumed to vary with synaptic area *s* ranging between 0 – 0.2 *μ*m^2^, as described by de Almeida et al. [27],

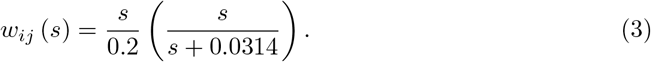

The probability density of the synaptic area of excitatory synapses onto granule cells in the model is given by [27]

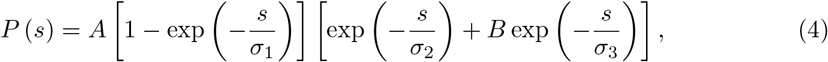

with *A* = 100.7, *B* = 0.002, *σ*_1_ = 0.022 *μ*m^2^, *σ*_2_ = 0.018 *μ*m^2^, and *σ*_3_ = 0.15 *μ*m^2^, as determined by Trommald and Hulleberg [120].

### Feedback inhibition

The effects of the diverse interneuron populations observed experimentally were approximated via one homogeneous population of ‘typical’ interneurons. Feedback inhibition was based on a model in which the first pyramidal cell to fire within a gamma-oscillation cycle excites an interneuron, which then inhibits the activity of later firing pyramidal neurons. In this rate-based model, it was assumed that neurons with greater excitatory input, and thus higher firing rates, are more likely to quickly overcome the decaying inhibition at the start of each gamma-oscillation cycle, and therefore fire earlier within each cycle [28]. Two modes of inhibition were employed, as specified: in the ‘winner-takes-all’ mode, only the most excited cell escaped inhibition, while in the ‘E%-max’ mode, cells were inhibited if their firing rate was not within some fraction of the excitation of the most excited cell [27]. E% was approximated to account for feedback delay, determined by the ratio of the membrane time constant to the time lag between a cell spiking and the incidence of feedback inhibition, which approximately gives E% = 10% [66].

The feedback inhibition mechanism initially compared pyramidal cell firing rates across the entire track. As specified in the Results section, the feedback inhibition cycle was shortened by incorporating position-specific inhibition within each bin along the track.

Thus, a pyramidal cell was inhibited when the condition

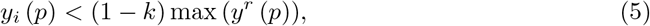

was satisfied, where *y_i_*(*p*) is the the firing rate of pyramidal cell *i* at position *p*, *k* determines the fraction (1 − *k*) of cells that escape inhibition, with *k* = 1 in the ‘winner-takes-all’ case and *k* = 0.1 in the ‘E%-max’ case, and *y^r^*(*p*) is the firing rates of all cells that project to interneuron *r*, at position *p*.

### Network architecture

The model was initially comprised of 10,000 grid cells and 2,000 pyramidal cells, with each pyramidal cell being innervated by 1,200 random grid cells, approximating the number of spatially-tuned EC cells that project to CA1 pyramidal cell, as described by [28]. The described feedback inhibition mechanism was mimicked by incorporating equal numbers of interneurons and pyramidal cells, with each interneuron receiving information from all pyramidal cells and projecting to a single cell each.

Where specified in the Results section, the network architecture was updated to fit experimental data on relative cell and synapse numbers in CA1 [59]. The network architecture was scaled to 5,000 grid cells, such that there were 7,788 pyramidal cells and 974 interneurons. Each pyramidal cell was innervated by 31 grid cells and 3 interneurons, with each interneuron being innervated by 19 pyramidal cells.

### Synaptic plasticity

#### Hebbian learning

Where specified, synaptic weights *w_ij_* between grid cell *j* and pyramidal cell *i* were updated once a day using Hebbian learning,

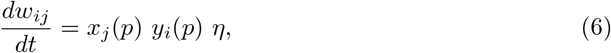

where *x_j_*(*p*) is the firing rate of grid cell *j* at position *p*, *y_i_* (*p*) is the firing rate of place cell *i* at position *p*, and *η* = 0.003 is the learning rate [28].

#### BCM learning

Where specified, synaptic weights were instead updated once a day according to the BCM rule [52],

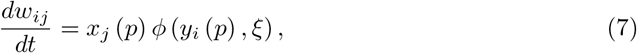

where 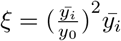 is the learning rate with positive constant *y*_0_ = 50 and *ϕ* (*y, ξ*) = *y* (*y − ξ*) with limit *l* = 2, such that

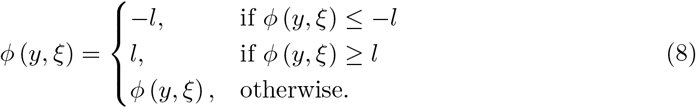

### Homeostatic mechanisms

Multiplicative scaling was applied to synaptic weights after each update by dividing by the sum of all weights converging onto each pyramidal cell and multiplying by the expected sum of synaptic strengths [121]. This value corresponded to the expected value of the sum of *n* random draws from the empirical distribution of weights, where *n* is the number of synapses converging onto each cell.

Where specified, subtractive normalization was instead applied after Hebbian learning [122]

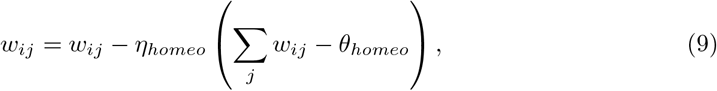

where *θ_homeo_* = 50 and *η_homeo_* is the homeostatic learning rate, set to 0.1 of the Hebbian learning rate to mimic the relative temporal timescale of Hebbian plasticity and homeostatic mechanisms in the hippocampus [78].

### Synapse turnover

Synapses were turned over once a day to achieve a mean synapse lifetime *τ* of 10 days [46]. The number of synapses replaced per cell was derived using the exponential decay model

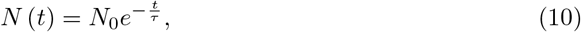

where *N* (*t*) and *N*_0_ are the number of synapses on day *t* and 0, respectively, and the number of synapses replaced per cell corresponds to *N*_0_ – *N* (*t*). Naive synapses received random weights from the synaptic strength distribution. In the scaled model, interneuron-to-pyramidal-cell connections were turned over at the same rate.

### AD-related synapse loss

Loss of grid-cell-to-pyramidal-cell synapses and interneuron-to-pyramidal-cell synapses were implemented individually and in combination, as specified, to analyze the effects of excitatory and inhibitory synapse loss on place cell function.

Interneuron-to-pyramidal-cell synapse loss due to axonal loss in the APP/PS1 model was implemented by eliminating a fraction of randomly chosen synapses each day using the quadratic regression for the change in the number of synapses Δ*S*(*T*) at timepoint *T* months from the first running session,

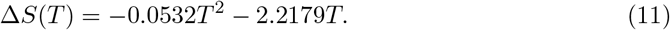

Grid-cell-to-pyramidal-cell synapses were eliminated according to the pattern of synaptic density reduction in APP/PS1 mice [67], using the regression

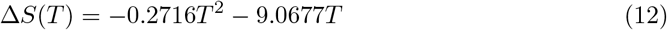

for *T ≤* 12 months.

### Place field analysis

A significant place field was defined as a single continuous region of 5 − 50 cm (see Results section), in which the cell’s firing rate was within 20% of the its maximum firing rate during the running session [28]. For calculating the probability of recurrence of a place cell, a place field was deemed to have been maintained across sessions if the location of the centroid was within 10 cm of the centroid in the previous session if samples were taken every 20 days, or 5 cm if samples were taken every 5 days.

## Acknowledgements

We acknowledge useful discussions with Dr. Mary Ann Go, Dr. Amanda Foust, Klara Kaleb and Grace Ang. This work was supported by Mrs Anne Uren and the Michael Uren Foundation.

